# Cerebello-cerebral Functional Connectivity Networks in Major Depressive Disorder: A CAN-BIND-1 Study Report

**DOI:** 10.1101/2021.06.25.449819

**Authors:** Sheeba Arnold Anteraper, Xavier Guell, Yoon Ji Lee, Jovicarole Raya, Ilya Demchenko, Nathan W. Churchill, Benicio N. Frey, Stefanie Hassel, Raymond W. Lam, Glenda M. MacQueen, Roumen Milev, Tom A. Schweizer, Stephen C. Strother, Susan Whitfield-Gabrieli, Sidney H. Kennedy, Venkat Bhat, the CAN-BIND Investigator Team

**Affiliations:** Department of Psychology, Northeastern University, Boston, MA, USA; Carle Foundation Hospital, Urbana, IL, USA; Alan and Lorraine Bressler Clinical and Research Program for Autism Spectrum Disorder, Massachusetts General Hospital, Boston, MA, USA; Department of Neurology, Massachusetts General Hospital and Harvard Medical School, Boston, MA, USA; Interventional Psychiatry Program, St. Michael’s Hospital, Toronto, ON, Canada; Keenan Research Centre for Biomedical Science, St. Michael’s Hospital, Toronto, ON, Canada; Neuroscience Research Program, St. Michael’s Hospital, Toronto, ON, Canada; Department of Psychiatry and Behavioural Neurosciences, McMaster University, Hamilton, ON, Canada; Mood Disorders Program and Women’s Health Concerns Clinic, St. Joseph’s Healthcare Hamilton, Hamilton, ON, Canada; Department of Psychiatry and the Mathison Centre for Mental Health Research and Education, University of Calgary, Calgary, AB, Canada; Department of Psychiatry, University of British Columbia, Vancouver, BC, Canada; Department of Psychiatry and Psychology, Queen’s University, Kingston, ON, Canada; Providence Care Hospital, Kingston, ON, Canada; Rotman Research Institute, Baycrest Health Sciences, Toronto, ON, Canada; Department of Biophysics, University of Toronto, ON, Canada; McGovern Institute for Brain Research, Department of Brain and Cognitive Sciences, Massachusetts Institute of Technology, Cambridge, MA, USA; Institute of Medical Science, University of Toronto, Toronto, ON, Canada; Department of Psychiatry, University of Toronto, ON, Canada

**Author notes:** **Corresponding Author:** Venkat Bhat, MD MSc, Interventional Psychiatry Program, St. Michael’s Hospital, 193 Yonge Street 6-012, Toronto, Ontario, Canada, M5B 1M4.

**Keywords:** Cerebellum, Default Mode Network, Depressive Disorder, Functional Magnetic Resonance Imaging, Biomarkers

## Abstract

**Objective:** Neuroimaging studies have demonstrated aberrant structure and function of the “cognitive-affective cerebellum” in Major Depressive Disorder (MDD), although the specific role of the cerebello-cerebral circuitry in this population remains largely uninvestigated. The objective of this study was to delineate the role of cerebellar functional networks in depression.

**Methods:** A total of 308 unmedicated participants completed resting-state functional magnetic resonance imaging scans, of which 247 (148 MDD; 99 Healthy Controls, HC) were suitable for this study. Seed-based resting-state functional connectivity (RsFc) analysis was performed using three cerebellar regions of interest (ROIs): ROI_1_ corresponded to default mode network (DMN) / inattentive processing; ROI_2_ corresponded to attentional networks including frontoparietal, dorsal attention, and ventral attention; ROI_3_ corresponded to motor processing. These ROIs were delineated based on prior functional gradient analyses of the cerebellum. A general linear model was used to perform within-group and between-group comparisons.

**Results:** In comparison to HC, participants with MDD displayed increased RsFc within the cerebello-cerebral DMN (ROI_1_) and significantly elevated RsFc between the cerebellar ROI_1_ and bilateral angular gyrus at a voxel threshold (*p* < 0.001, two-tailed) and at a cluster level (*p* < 0.05, FDR-corrected). Group differences were non-significant for ROI_2_ and ROI_3_.

**Conclusions:** These results contribute to the development of a systems neuroscience approach to the diagnosis and treatment of MDD. Specifically, our findings confirm previously reported associations between MDD, DMN, and cerebellum, and highlight the promising role of these functional and anatomical locations for the development of novel imaging-based biomarkers and targets for neuromodulation therapies.

## Introduction

Psychiatric disorders are highly prevalent and account for almost one quarter of the global burden of disease [1]. The greatest contributor is Major Depressive Disorder (MDD), the prevalence of which has increased significantly since 1990 [2]. Current estimates suggest that about 350 million people across the globe suffer from MDD. Its annual economic burden is approximated to be in the range of several hundred billion dollars, with 50% attributable to workplace costs, 45% to direct costs, and 5% to suicide-related costs [3]. As a heterogeneous disorder with diverse and poorly understood etiology, the treatment of MDD is particularly challenging. Only one-third or less of patients will achieve remission on an initial trial of antidepressant medication, while another one-third will remit after trying other antidepressant and augmentation strategies [4]. Even when antidepressants are effective, there is often a time delay of several weeks before response or remission is achieved [5]. There is a strong need to improve our mechanistic understanding of the pathogenesis and disease progression of MDD so that novel therapeutic strategies can be developed.

Over the past decade, structural and functional neuroimaging studies have characterized the non-motor functions of the cerebellum and its involvement in cognitive and affective processing [6–8]. Lesions to the vermis and the posterior lobe frequently induce the cerebellar cognitive affective syndrome – a condition characterized by emotional lability, impaired social cognition, depressed mood, and deficits in executive functioning and linguistic processing [9]. Several functional magnetic resonance imaging (fMRI) studies have confirmed engagement of the non-motor cerebellum in intrinsic connectivity networks (ICNs), including the cognitive control network (CCN) [10, 11], the salience network (SN) [10, 12], the default mode network (DMN) [12, 13], and the cerebello-amygdaloid network [14]. In fact, resting-state functional connectivity (RsFc) analysis suggests that approximately half of the cerebellar cortex comprises the “cognitive-affective cerebellum”, where non-motor functions are represented within lobules VI-IX of the posterior lobe [6].

Cognitive dysfunction is a prominent symptom during a Major Depressive Episode (MDE) and is often mediated by affective dysregulation [15]. Core diagnostic criteria include diminished ability to think, lack of concentration, distractibility, and indecisiveness. Mood disturbances further interact with the cognitive symptom dimension, manifested in distorted information processing, negativity bias, and abnormal response to negative feedback [16].

Based on RsFc studies, MDD patients exhibit reduced connectivity of the cerebellar lobule VIIA and crus I/II with cortical components of the CCN - the dorsolateral prefrontal cortex (dlPFC) [17, 18] and the ventromedial prefrontal cortex (vmPFC) [17, 19]. Aberrant lobule VIIA-vmPFC coupling has been further shown to be associated with impaired verbal working memory performance [17] and the antidepressant efficacy of electroconvulsive therapy [20]. Acting as a modulator within the SN, the vermis of lobule VI is thought to represent a phylogenetically older emotional processor [12]. Structurally, its volume is positively correlated with depressive symptoms, mostly along the somatic symptom subscale [21]. Within the DMN, the key findings highlight reduced functional connectivity between lobule VII and crus I with the inferior parietal and temporal cortices [22] as well as altered connectivity of the vermis with the anterior cingulate cortex (ACC) and the vmPFC [19]. Evidence concerning lobule VIIA-posterior parietal cortex (PCC) functional connectivity is less consistent, as both increased [17] and decreased [18] coupling have been reported. Moreover, there is anatomical evidence suggesting that lobule IX participating in the DMN is bigger in volume in patients with both acute and remitted MDD [12, 23].

Although cerebello-cerebral connectivity deficits have been demonstrated to contribute to MDD brain profile, the specific role of the cerebellar circuitry in depression remains largely unexplored. A recent RsFc analysis of the dataset from the initial trial of the Canadian Biomarker Integration Network in Depression (CAN-BIND-1) revealed greater connectivity between the insula seed (part of the SN) and cerebellar lobules VI, VIIA (crus I & II), and VIII among participants with MDD who completed a course of antidepressant medication therapy [24]. Notably, that insular-cerebellar connectivity pattern differentiated remitters from non-remitters and showed the involvement of posterior cerebellar regions associated with affective processing. Using the same dataset, the objective of the current study was to compare RsFc in participants in an MDE with MDD and healthy controls (HC) to test for depression-associated connectivity differences between the functionally distinct regions of the cerebellar cortex and the cerebral cortex. Contrary to previous investigations, the whole-brain seed-to-voxel RsFc analysis performed in this study had a cerebellar focus with the seed regions of interest (ROIs) derived from functional gradient analyses of the cerebellum [25].

## Methods

### Study Procedure and Participants

We analyzed the resting-state functional and structural MRI data from the CAN-BIND-1 trial (ClinicalTrials.gov identifier: NCT01655706; Registration date: August 2^nd^, 2012) [26, 27]. Full details of the CAN-BIND-1 trial have been described elsewhere [26, 27]. Participants diagnosed with MDE in context of MDD (*N* = 200) and matched healthy controls (*N* = 108) were recruited at 6 Canadian clinical centres: University Health Network (Toronto Western/Toronto General Hospital) in Toronto, Ontario; Centre for Addiction and Mental Health in Toronto, Ontario; St. Joseph’s Healthcare Hamilton in Hamilton, Ontario; Providence Care Hospital in Kingston, Ontario; Djavad Mowafaghian Centre for Brain Health in Vancouver, British Columbia; and Hotchkiss Brain Institute in Calgary, Alberta. Standard participation and data transfer agreements were regulated by the Ontario Brain Institute (https://www.braincode.ca/content/governance) as well as local ethical and legislative bodies at each institution. All participants provided written informed consent at the Screening Visit. Participants in the depression group were adult outpatients between 18-60 years of age with a primary DSM-IV-TR diagnosis of MDD, as established from the Mini-International Neuropsychiatric Interview [28], and had symptom scores of of ≥ 24 on the Montgomery-Åsberg Depression Rating Scale (MADRS). Detailed inclusion and exclusion criteria for both groups of participants are outlined elsewhere [26]. The final samples suitable for the analyses of cerebello-cerebral RsFc were 148 MDD patients and 99 HC. From the MDD group, 52 participants were excluded due to missing clinical data (N = 23), non-adherence to CAN-BIND-1 protocol (N = 10), and missing fMRI data or poor fMRI data quality (N = 19). From the control group, 9 participants were excluded due to poor fMRI data quality.

### Selection of cerebellar regions of interest

Three ROIs were selected based on the two principal gradients configuring the gradual organization of cerebellar functional regions: from motor to task-unfocused regions (*gradient 1*), and from task-focused to task-unfocused regions (*gradient 2*) (**Figure 1**), as described previously by Guell and colleagues [25]. Binary masks of voxels from gradient extremes defined ROI_1_ (gradient 1, top 5% values) as corresponding to inattentive / DMN processing; ROI_2_ (gradient 2, top 5% values) corresponding to attentional networks including frontoparietal network (FPN), dorsal attention network (DAN), ventral attention network (VAN); and ROI_3_ (gradient 2, bottom 5% values) corresponding to motor processing. Readers are referred to the original description of functional gradients of the cerebellum by Guell and colleagues [25] for further methodological and theoretical background of cerebellar functional gradients; specifically see supplementary materials in elifesciences.org/articles/36652#fig1s1 for a description of the methodology of functional gradients, and elifesciences.org/articles/36652#fig3s3 for a description of the evidence that supports the functional significance of each ROI used in our analyses.

**Fig. 1.**
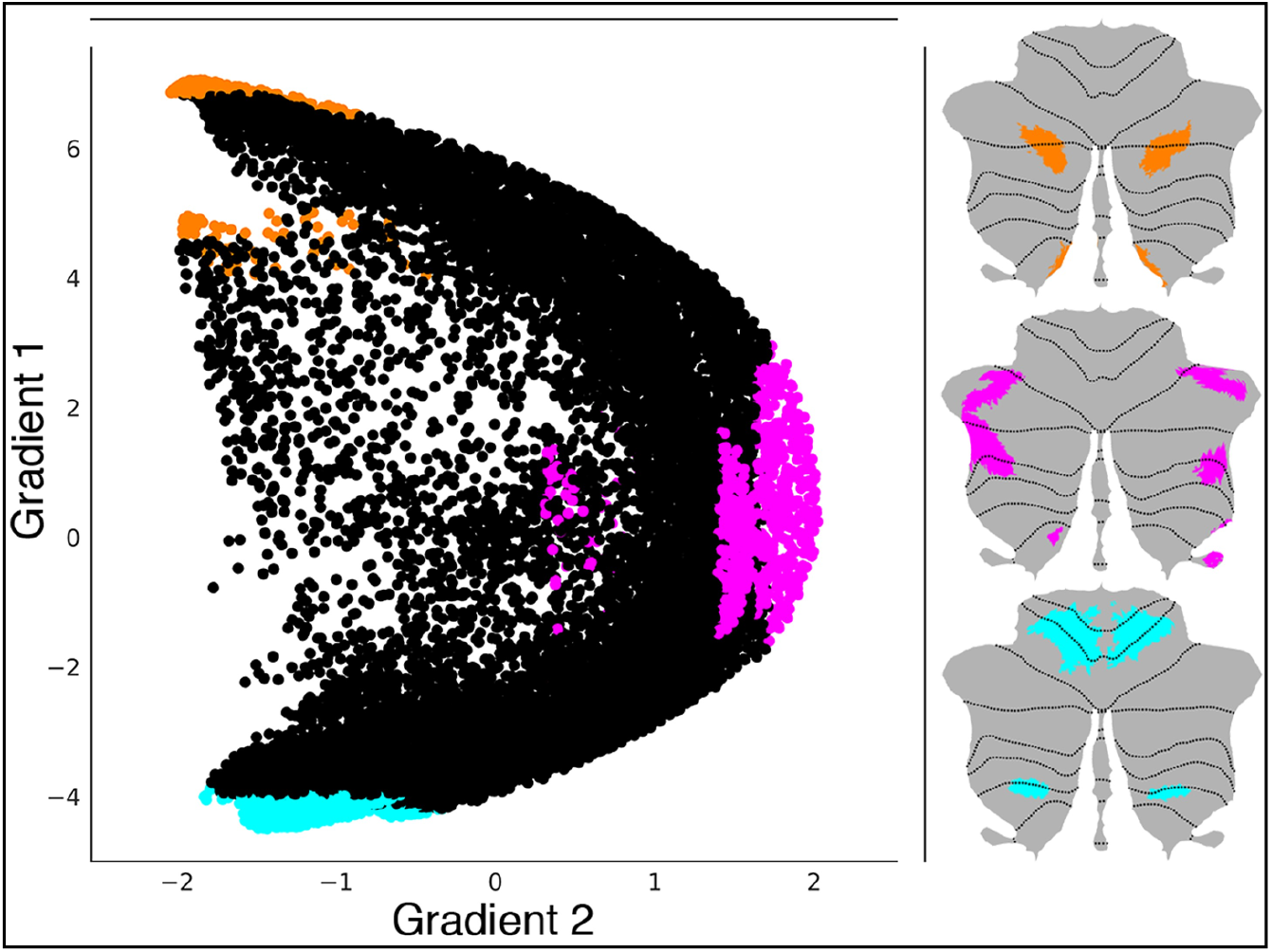
Cerebellar ROIs for whole-brain seed-to-voxel analysis were selected based on the functional gradients of the cerebellum [25]. Three maps were generated to locate gradient extremes at each area of motor and non-motor representation. Orange, pink and cyan colors depict gradient 1 top 5% values within each area of non-motor representation (VI/Crus I, contiguous Crus II/VIIB, and IX, X), gradient 2 – the top 5% values within each area of non-motor representation (VI/Crus I, Crus II/VIIB, and IX, X), and gradient 1 – the bottom 5% values within each area of motor representation (I-VI and VIII), respectively

### Neuroimaging Data Collection

High-resolution (1 mm^3^ isotropic) 3-dimensional isotropic T1-weighted scans and resting-state fMRI data were collected at all sites according to the protocol published elsewhere [27]. The resting-state fMRI scans were acquired using a whole-brain T2*-weighted blood-oxygen-level-dependent echo planar imaging sequence. The following scanner models were used for data collection: Signa HDxt 3.0 T (GE Healthcare), Discovery MR750 3.0 T (GE Healthcare), MAGNETOM Trio (Siemens Healthcare), and Intera 3.0 T (Philips Healthcare). Acquisition parameters are provided in Table 1. Instructions and support materials were standardized across sites [27].

**Table 1.** Resting-state functional MRI datasets (duration = 10 minutes) and acquisition parameters.

| CAN-BIND-1 dataset | Toronto<br>Western/Toronto<br>General Hospital | Centre for<br>Addiction and<br>Mental Health | St. Joseph's<br>Healthcare<br>Hamilton | University of<br>Calgary | University of<br>British Columbia | Queen's<br>University |
| --- | --- | --- | --- | --- | --- | --- |
| Scanner model | GE 3.0 T<br>Signa HDxt | GE 3.0 T<br>Discovery MR750 | GE 3.0 T<br>Discovery MR750 | GE 3.0 T<br>Discovery MR750 | Phillips 3.0 T<br>Intera | Siemens 3.0 T<br>TrioTim |
| Repetition time (ms) | 2000.0 | 2000.0 | 2000.0 | 2000.0 | 2000.0 | 2000.0 |
| Echo time (ms) | 30.0 | 30.0 | 30.0 | 30.0 | 30.0 | 30.0<sup>a</sup> |
| Flip angle (degrees) | 75.00 | 75.00 | 75.00 | 75.00 | 75.00<sup>b</sup> | 75.00 |
| Slices (*n*) | 34<sup>c</sup> | 36<sup>d</sup> | 36 | 36 | 36 | 36<sup>e</sup> |
| Voxel resolution (mm<sup>3</sup>) | 4 x 4 x 4 | 4 x 4 x 4 | 4 x 4 x 4 | 4 x 4 x 4 | 4 x 4 x 4 | 4 x 4 x 4 |

CAN-BIND = Canadian Biomarker Integration Network in Depression.
<sup>a</sup>For *n* = 11, echo time = 25.0 ms.
<sup>b</sup>For *n* = 27, flip angle = 90.00 degrees.
<sup>c</sup>For *n* = 40, number of slices = 40.
<sup>d</sup>For *n* = 7, number of slices = 40.
<sup>e</sup>For *n* = 11, number of slices = 40.
Detailed scan acquisition parameters for resting-state functional MRI sequences are available elsewhere [27].

### Data Processing: First-level Seed-to-voxel Functional Connectivity Analysis

Resting-state MRI scans were realigned and spatially normalized to the MNI template (4th Degree B-Spline) and smoothed with an isotropic 6 mm FWHM Gaussian kernel using SPM12 (Wellcome Department of Imaging Neuroscience; www.fil.ion.ucl.ac.uk/spm). T1-weighted anatomical scans were brought to the MNI space and then partitioned into white matter (WM), gray matter, and cerebrospinal fluid (CSF) using SPM’s tissue priors employing the automatic segmentation routine in SPM12. The CONN Toolbox [29] was used to carry out denoising steps: band-pass filtering (0.008-0.1 Hz) and removal of physiological confounds (to eliminate contributions from WM and CSF [30]). Motion outliers can increase the likelihood of Type II errors (decreasing sensitivity) as well as Type I errors (affecting the validity of the analyses). Therefore, we used a stringent scan-to-scan threshold of <0.5 mm composite motion, and <3 standard deviations of global mean signal to identify outliers using the CONN Toolbox [37] and as part of the denoising step, motion outliers and their first-order derivatives along with motion outliers were regressed out. Whole-brain correlation maps from the seed ROIs were then computed. ROIs included voxels from the three gradient extremes [25, 31]: specifically, the top 5% values of gradient 1 (corresponding to DMN and language), the top 5% values of gradient 2 (corresponding to FPN, DAN, VAN, and working memory), and the bottom 5% values of gradient 1 (corresponding to motor processing) (see **Figure 1**). For the first-level seed-based RsFc analysis, Pearson’s correlation coefficients were generated by computing the correlations between the time series of the ROIs and time series of the rest of the voxels in the brain volume.

### Data Processing: Second-Level General Linear Model (GLM) Analysis

Whole-brain seed-to-voxel *r*-maps from the first-level were then transformed to *z*-maps (Fisher’s *r*-to-*z* transform) and voxelwise GLM analysis was conducted on connectivity values at the second level for within-group and between-group comparisons. Statistical significance thresholding for between-group effects were *p* < 0.001 (two-tailed) at the voxel level and *p* < 0.05 False Discovery Rate (FDR) correction at the cluster level [32].

## Results

### Sociodemographic and Clinical characteristics

The clinical characteristics of the sample included in this study are provided in Table 2. Age, sex, head motion parameters, and outliers of motion were not statistically different between groups.

**Table 2.** Baseline sociodemographic and clinical characteristics of MDD patients and Healthy Controls.

| | MDD, $N = 148$ | HC, $N = 99$ | $p$-value |
| --- | --- | --- | --- |
| **Sociodemographic characteristics** |  |  |  |
| Sex, $N$ male (%) | 57 (38.51) | 35 (35.35) | .712 |
| Age, mean ($SD$) years | 34.43 (12.35) | 33.02 (10.87) | .357 |
| Education, mean ($SD$) years | 16.86 (2.13) | 18.32 (2.31) | < .001 |
| Ethnicity, $N$ white (%) | 113 (76.35) | 70 (70.71) | .321 |
| Marital status, $N$ nonmarried<sup>a</sup> (%) | 106 (71.62) | 65 (65.66) | .393 |
| Employment, $N$ working/student (%) | 95 (64.19) | 91 (91.92) | < .001 |
| Handedness, $N$ right-handed (%) | 130 (87.84) | 85 (85.86) | .794 |
| **Clinical characteristics** |  |  |  |
| MADRS Total, mean ($SD$) | 29.74 (5.64) | 0.90 (1.75) | < .001 |
| Age of MDD onset, mean ($SD$) | 19.48 (9.13) | - | - |
| Current MDE duration, $N$ more than 1 year (%) | 67 (45.27) | - | - |
| Current MDE duration in months, mean ($SD$) | 27.35 (34.77) | - | - |
| Psychiatric comorbidities<sup>b</sup> |  |  | - |
| Anxiety disorders<sup>c</sup>, $N$ (%) | 67 (45.27) | 0 (0.0) | - |
| Substance-related disorders<sup>d</sup>, $N$ (%) | 7 (4.73) | 0 (0.0) | - |
| Eating disorders<sup>e</sup>, $N$ (%) | 2 (1.35) | 0 (0.0) | - |

HC = Healthy Controls; MADRS = Montgomery-Åsberg Depression Rating Scale; MDD = Major Depressive Disorder; MDE = Major Depressive Episode.
<sup>a</sup>Includes single, divorced, widowed, and separated.
<sup>b</sup>Based on the DSM-IV-TR, as determined by the Mini-International Neuropsychiatric Interview. Patients may have more than one psychiatric comorbidity.
<sup>c</sup>Includes panic disorder, agoraphobia, social phobia, obsessive-compulsive disorder, posttraumatic stress disorder, and generalized anxiety disorder.
<sup>d</sup>Includes alcohol abuse and substance use.
<sup>e</sup>Includes anorexia nervosa and bulimia nervosa.

### Second Level GLM Analysis

RsFc analysis from the cerebellar ROI_1_ (top 5% values of gradient 1) revealed increased cerebello-cerebral connectivity in the MDD group compared to HC, within the brain areas corresponding to DMN (the precuneus (PCu), the posterior cingulate cortex (PCC), the medial prefrontal cortex (mPFC), and the inferior parietal cortices (IPL) (**Figure 2**, black arrows)). Of these regions, the bilateral angular gyrus (AG, peak coordinates at [−38, −56, 32] and [58, −56, 32]) was significant at a voxel threshold of *p* < 0.001 (two-tailed) and *p* < 0.05 (FDR-corrected) at the cluster level. Violin plots of the significant clusters are provided in **Figure 3**. Second-level RsFc analyses from the other two ROIs (ROI_2_, top 5% values of gradient 2; ROI_3_, bottom 5% values of gradient 1) did not reveal any statistically significant differences between MDD and HC participants.

**Fig. 2.**
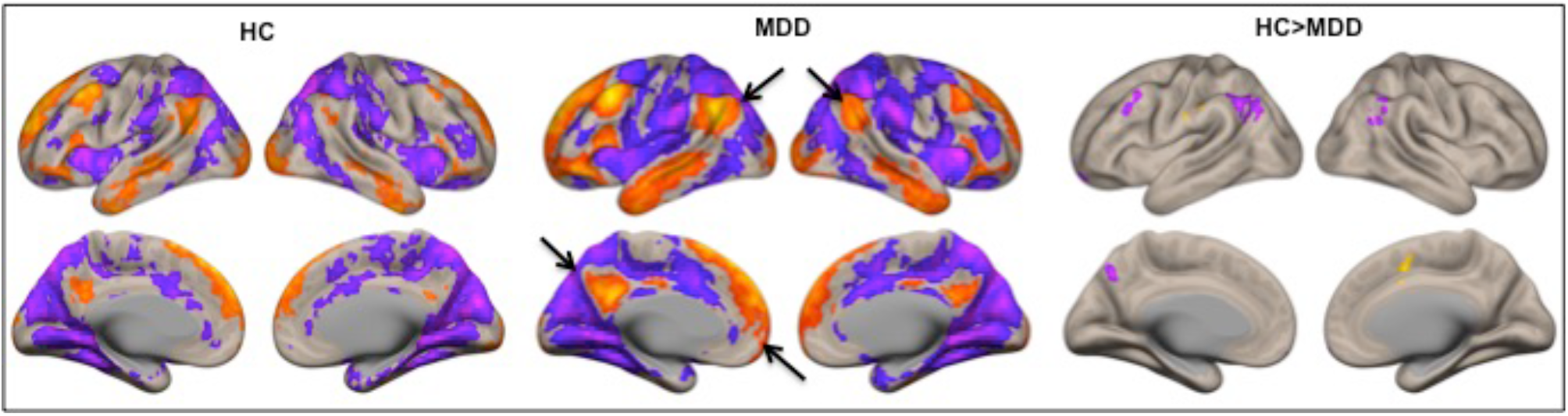
Within-group (HC, MDD) and between-group (HC versus MDD) RsFc results for the top 5% values of the cerebellar gradient 1 (height-threshold of p < 0.005, two-tailed; p < 0.05, FDR-corrected). Clusters pointed by black arrows correspond to the posterior cingulate cortex (PCC), the medial prefrontal cortex (mPFC) and the inferior parietal cortices (IPL), which overlap with the default mode network (DMN) [33]. Compared to HC, the MDD group displayed a significantly greater (height-threshold of p < 0.001, two-tailed; p < 0.05, FDR-corrected) functional connectivity between the cerebellar ROI_1_ and the bilateral angular gyrus (AG). Positive correlations are shown in yellow and negative correlations in purple

**Fig. 3.**
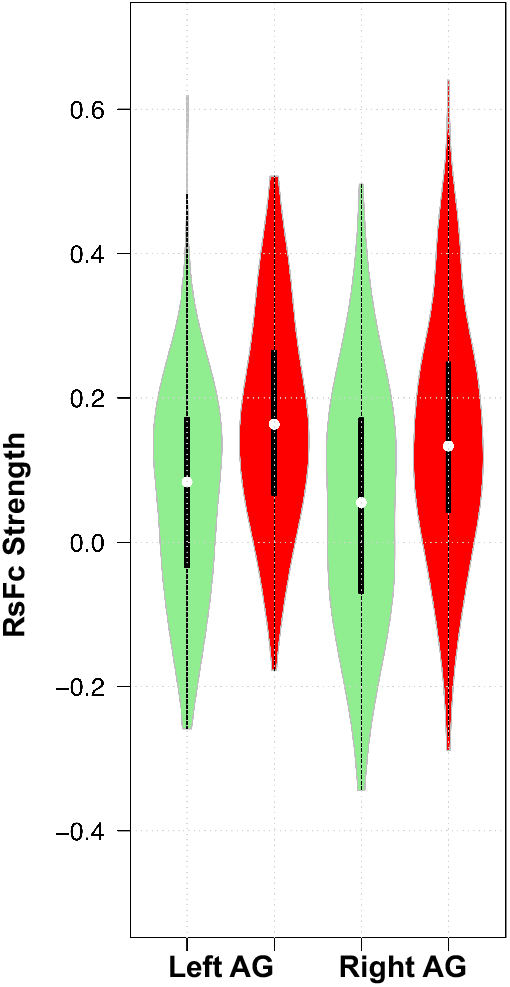
Violin plots for significant clusters from RsFc analysis of cerebellar ROI_1_ in the HC (green) and MDD (red) participant groups. White circles show the medians (MDD, left angular gyrus: 0.16, right angular gyrus: 0.13; HC, left angular gyrus: 0.08, right angular gyrus: 0.06); box limits indicate the 25th and 75th percentiles as determined by the R software; whiskers extend 1.5 times the interquartile range from the 25th and 75th percentiles; polygons represent density estimates of data and extend to extreme values

## Discussion

We used data-driven clustering to segment the cerebellum and examine the RsFc of cerebellocortical networks in MDD. Of the three cerebellar ROIs explored, we report abnormal connectivity within the cerebello-cerebral DMN in the MDD group compared to controls. MDD participants displayed a significantly higher RsFc between the cerebellar ROI_1_ (lobules VI/Crus I, contiguous Crus II/VIIB and IX/X) and the bilateral AG of the cerebral cortex. This evidence illustrates the use of the functional gradient approach as a promising method to examine network-level changes associated with psychiatric disorders and exemplifies the potential use of functional connectivity between cerebellar non-motor regions and the cerebral cortex as a potential underlying mechanism involved in cognitive and affective processing in depression.

### Cerebello-cerebral DMN in Depression

The lobules of the “cognitive-affective cerebellum” have complex topographic organization, consisting of subregions with distinct functions and connectivity patterns [7]. Based on existing neuroimaging evidence, the non-motor cerebellum contributes to multiple cortical ICNs implicated in depression, including the CCN, DMN, SN, and CAN. The DMN, one of the most commonly studied ICNs, plays a pivotal role in the neurobiology of MDD, specifically for symptoms characterized and explained by disturbances in emotional modulation, self-referential activity, episodic memory retrieval, and intrinsic attention allocation [34]. Previous research has investigated the role of the cerebellum in the aberrant activity of the DMN in depression, although results have been mixed. In patients with MDD, relative to HCs, both increased [17] and decreased [18, 22, 35] cerebello-cerebral functional connectivity have been reported. Studies of whole-brain network organization suggest that potentially recruited cerebellar subregions include the posterior part of lobule VIIA, crus I, crus II, and, potentially, lobule IX [6, 36, 37]. It should be noted, however, that these previous studies did not consider functional parcellation of the cerebellum based on cerebellar functional gradients, which capture the organization of the cerebellum in a hierarchical way [32]. Our study is the first to employ this methodology for evaluating cerebello-cerebral connectivity differences in MDD.

The DMN is comprised of the PCu, the PCC, the mPFC, rostral ACC, bilateral IPL, and medial/lateral temporal cortices [33]. When healthy individuals engage in self-referential thinking, DMN connectivity increases, serving as a measure of internally generated cognition [38]. In contrast, DMN deactivation is seen upon task switching or increasing task difficulty, possibly reflecting a reallocation of resources from resting-state toward attentional and cognitive control processes [33]. Resting-state fMRI studies provide substantial evidence that patients with depression exhibit DMN hyperactivity during self-referential processes, which does not return to baseline levels during attention-demanding tasks [39] or a shift from intrinsic to externally focused thoughts [40]. Similar to the degree of cerebellar involvement in fine-tuning motor activity and regulating balance and posture, certain areas have a critical role in modulating fine aspects of human cognition and emotions, thereby acting as an “emotional pacemaker” [41]. Relative hyperconnectivity with cerebral regions possibly originates from the loss of normal modulation of the DMN circuitry, leading to the generation of more extreme swings within internally-generated cognitive processes and overcorrections – a state somewhat analogous to dysmetria diagnosed from a finger-to-nose or a heel-to-shin test [42].

The AG (Brodmann area 39), which resides in the posterior part of the IPL, is involved in a wide array of cognitive functions, including theory of mind, social cognition, semantic processing and language, conceptual representation, visuospatial navigation, and episodic memory [43]. As a region with widespread structural and functional connections, particularly to other densely-connected regions of the brain, it not only constitutes one of the core DMN-associated areas, but also drives communication within other ICNs, notably the FPN and the VAN [44]. The AG emerges as a “connector hub”, i.e., a region that receives multisensory information from functionally distinct regions of the brain, including those implicated in attention allocation and emotion regulation [43, 45]. In turn, this likely facilitates integration of complex mental representations and interpersonal information in social situations, highlighting the functional role of bilateral AG as dynamic brain areas exerting a top-down influence on self-referential processes [46]. An emerging body of evidence supports the role of aberrant structural and functional AG connectivity in depression [47–50]. As part of the DMN, the AG was proposed to serve as a central cross-modal hub of a deficient theory of mind subnetwork, linked to MDD-associated impairments in integrating contextual information for reasoning in social situations, extensive processing of self-referential information, and emotion dysregulation [47]. Moreover, as part of the DMN, the AG may be associated with rumination [51] and the adjustment of negativity bias [52]. Here, we report an increased functional connectivity between bilateral AG and the cerebellar ROI_1_ (lobules VI/Crus I, contiguous Crus II/VIIB and IX/X) in depression – a novel finding that expands understanding of the DMN connectivity profile and highlights the role of the cerebellum in self-referential processing, theory of mind, and rumination.

Several recent studies have focused on the role of the cerebellum in social cognition and theory of mind, with evidence largely derived from lesion and fMRI studies (for review, see [53]). Individuals with cerebellar damage show impaired performance in theory of mind, perceptual, mental state inference, and cognitive tasks, as well as in tasks involving social emotions [54]. Cerebellar abnormalities and dysfunction of the “social” cerebello-cerebral networks have previously been associated with characteristic mentalizing impairments, notably among patients with autism [55] and schizophrenia spectrum disorders [56]. Based on the observed hyperconnectivity between the cerebellar cortex and the bilateral AG, our study emphasizes the role of the “social cerebellum” in the pathophysiology and symptomatology of MDD. Similar to how the cerebellum operates within the sensorimotor domain, it most likely sends information about prediction errors to the DMN-associated areas, including the AG, triggering comparison with cross-modal afferent inputs received from other ICNs. In the presence of any deviations, the cerebellum would correct the signal, thereby updating a pre-generated representation within the internal socioemotional system. Our results indicate that MDD likely disturbs the intrinsic dynamics of this system leading to overcorrections, which might possibly translate into rumination and theory of mind deficits.

Within the cerebral cortex, individuals with MDD have increased levels of self-referential representations associated with higher levels of maladaptive, depressive rumination and lower levels of adaptive, reflective rumination [57]. Rumination, which is a central aspect of the phenomenology of MDD, refers to a tendency to perseverate about one’s own symptoms and often occurs when top-down control mechanisms facilitate patterns of negative self-referential processing and maintain a heightened focus on negative emotional states [58]. Enhanced within-DMN connectivity has been consistently associated with depressive rumination [39, 51]. Additionally, hyperconnectivity of the DMN in brain areas associated with affective processing and salient stimuli, such as the subgenual prefrontal cortex and the ACC, may serve as a primary pathophysiological factor contributing to depressive rumination [51]. The growing literature on the DMN supports the notion that clinically depressed patients show increased functional connectivity within this network [59], which can be at least partially normalized by antidepressants [60]. Our results provide further evidence of a strong association between DMN functional abnormalities and MDD and highlight the importance of the cerebellum in ICN theories of depression.

### Other Cerebello-cerebral Intrinsic Connectivity Networks in Depression

Our study did not reveal significant functional connectivity between the cerebellum and the ROI_2_ (VI/Crus I, Crus II/VIIB, and IX, X), corresponding to attentional networks including FPN, DAN, and VAN, or the ROI_3_ (I-VI and VIII), corresponding to motor representation. Negative results in fMRI research are challenging to interpret. When analyzing the signal in a given network of interest, the lack of significant differences between two groups does not prove the absence of functional alterations in these territories. However, the fact that significant differences were detected in the DMN (ROI_1_) but not in attentional (ROI_2_) or motor networks (ROI_3_) indicates that functional abnormalities, as indexed by the fMRI, may be more prominent in the DMN compared to these other networks in depression, as suggested by prior investigations reviewed earlier in this manuscript. Future studies aimed at directly comparing the degree of abnormalities between networks in MDD patients are required to confirm this possibility, since the finding that one network is significantly different between two groups while others are not, does not necessarily equate to differences between the abnormalities in these two networks being statistically significant.

### Limitations

Our study is not without limitations. Firstly, the neuroimaging findings presented here are based on exploratory analyses demonstrating the presence of functional alterations within the cerebello-cerebral network in MDD, with no direction of connectivity specified. Based on our understanding of anatomical connections between the cerebellum and the cerebral cortex, causal modeling studies are needed to examine the effective connectivity within the network and characterize the directionality of communication between the cerebellum and the cerebral cortex. Further, while the 3-cluster approach employed in this study is good for robustness, more exploration of finer subdivisions of the cerebellum in the future might yield improved specificity. Secondly, our results are limited to the MRI acquisition parameters employed across CAN-BIND-1 trial sites. Since those scanners a priori provide better spatial and temporal resolution of the cerebral cortex, the MDD-associated cerebello-cerebral hyperconnectivity that we report here needs to be replicated and validated using alternative MRI acquisition protocols and scanners of ultra-high field strength (e.g., 7 T). Given the heterogeneity of MDD, future work could also examine MDD cohort variability and possible demographic drivers, as our results are limited to a specific sample of participants with MDD.

### Conclusion

This study demonstrates that patients with MDD exhibit abnormal functional cerebello-cerebral hyperconnectivity between the cerebellum and the AG within the DMN, which might contribute to the cognitive-affective dysfunction characteristic of depression. Studying cerebellar networks and the hindbrain, phylogenetically the oldest brain structures, will help delve into the pathophysiology of depression from previously uncharted angles, possibly identifying important network constituents carrying integral modulatory function. The findings of our study advance the overall mechanistic understanding of the role of cerebellum in non-motor functions traditionally researched in the context of neocortical higher order processing. Our results, additionally, support numerous studies that have previously reported aberrant activity of the DMN in depression, thus contributing to the vast literature on ICNs in affective disorders. Finally, cerebello-cerebral functional connectivity might be a potential biomarker of depression in clinical and research practices, where the DMN regions of cerebellum may be employed as potential targets for novel non-invasive therapeutic interventions.

## Acknowledgements

The authors would like to acknowledge the contributions of Mojdeh Zamyadi and Jacqueline Harris for data quality control, of Andrew Davis and Geoffrey Hall for sequence assessment and standardization, and of Yuelee (Ben) Khoo for statistical analysis of the demographics data.

## Author Contributions

Study conceptualization: Sheeba Arnold Anteraper, Venkat Bhat; Methodology development: Sheeba Arnold Anteraper, Nathan. W. Churchill, Venkat Bhat; Formal analysis and investigation: Nathan W. Churchill, Tom A. Schweizer, Venkat Bhat; Writing – original draft preparation: Sheeba Arnold Anteraper, Xavier Guell, Ilya Demchenko, Venkat Bhat; Writing – review and editing: Sheeba Arnold Anteraper, Xavier Guell, Yoon Ji Lee, Jovicarole Raya, Ilya Demchenko, Nathan W. Churchill, Benicio N. Frey, Stefanie Hassel, Raymond W. Lam, Glenda M. MacQueen, Roumen Milev, Tom A. Schweizer, Stephen C. Strother, Susan Whitfield-Gabrieli, Sidney H. Kennedy, Venkat Bhat. All authors read and approved the final version of the manuscript. The data acquisition was supported by the CAN-BIND Investigator Team. There were no separate resources for this project.

## Compliance with Ethical Standards

### Ethics Approval

The authors assert that all procedures contributing to this work comply with the ethical standards of the relevant national and institutional committees on human experimentation and with the Helsinki Declaration of 1975, as revised in 2008. All study procedures were approved by the Institutional Ethics Board at University Health Network, Toronto, Ontario, Canada; Centre for Addiction and Mental Health, Toronto, Ontario, Canada; St. Joseph’s Healthcare, Hamilton, Ontario, Canada; Providence Care Hospital, Kingston, Ontario, Canada; Djavad Mowafaghian Centre for Brain Health, Vancouver, British Columbia, Canada; and Hotchkiss Brain Institute, Calgary, Alberta, Canada.

### Consent to Participate and Publish

All participants provided written, informed consent to participate in the study and have their data published.

